# PacRAT: A program to improve barcode-variant mapping from PacBio long reads using multiple sequence alignment

**DOI:** 10.1101/2021.11.06.467314

**Authors:** Chiann-Ling C Yeh, Clara J Amorosi, Soyeon Showman, Maitreya J Dunham

**Author notes:** Co-first authors (Chiann-Ling C Yeh, Clara J Amorosi).

## Abstract

**Motivation:** Use of PacBio sequencing for characterizing barcoded libraries of genetic variants is on the rise. PacBio sequencing is useful in linking variant alleles in a library with their associated barcode tag. However, current approaches in resolving PacBio sequencing artifacts can result in a high number of incorrectly identified or unusable reads.

**Results:** We developed a PacBio Read Alignment Tool (PacRAT) that improves the accuracy of barcode-variant mapping through several steps of read alignment and consensus calling. To quantify the performance of our approach, we simulated PacBio reads from eight variant libraries of various lengths and showed that PacRAT improves the accuracy in pairing barcodes and variants across these libraries. Analysis of real (non-simulated) libraries also showed an increase in the number of reads that can be used for downstream analyses when using PacRAT.

**Availability:** PacRAT is written in Python and is freely available on GitHub (https://github.com/dunhamlab/PacRAT).

## Introduction

Improvements in sequencing technology have greatly expanded our capacity not just for detecting genetic variants, but also for assessing their function. Approaches such as deep mutational scanning (DMS) use a library of variants and have been used to characterize the impact of genetic variation on protein structure, gene expression, and gene function across many fields (Matreyek *et al*., 2018; Thompson *et al*., 2020; Hoggard *et al*., 2018; Starita *et al*., 2015; Rich *et al*., 2016). Such variant libraries are commonly “barcoded” with short random DNA tags, which allows for short-read sequencing of multiple time points and removes the step of directly sequencing entire variants at every time point. Barcodes are associated with library variants, which is a simple process if the variable region and the linked barcode can be sequenced together with short-read methods. However, with longer variable regions this becomes increasingly costly, challenging, and time-consuming (Hiatt *et al*., 2010). Many studies are turning to PacBio sequencing to generate high-quality long reads that span the entirety of the barcode and variable region (Matreyek *et al*., 2018; Suiter *et al*., 2020; Ollodart *et al*., 2021; Amorosi *et al*., 2021; Yeh *et al*., 2021).

Compared to Illumina short-read sequencing, PacBio generates much longer continuous reads, though with the disadvantage of an increased error rate, which can be as high as 15% (Wenger *et al*., 2019). To reduce errors, highly accurate reads are generated from circular consensus sequencing (CCS), which circularizes molecules by blunt ligating single-stranded SMRTbell™ adaptors and allowing a single DNA molecule to be sequenced multiple times. Each pass through the molecule produces a subread, and these subreads are collapsed to form a consensus sequence that improves the quality score of a sequence (Rhoads and Au, 2015; Wenger *et al*., 2019). Although PacBio chemistry has improved, errors are still pervasive in CCS reads. Compared to Illumina NovaSeq, the PacBio mismatch rate is 17X lower, but indel rates are 181X higher (Wenger *et al*., 2019). Such indel errors are more prominent in CCS reads with lower numbers of subreads, often those from longer target sequences. In a recent study that performed PacBio barcode-variant mapping, more than 30% of barcodes were associated with indels and were discarded during analysis due to variation in length (Matreyek *et al*., 2018). Some libraries include variants that have real indels, which cannot always be ignored or discarded. Therefore, additional error correction steps are required to improve reliability and maximize information gained from PacBio consensus reads.

In this paper, we present **Pac**Bio **R**ead **A**lignment **T**ool (“PacRAT”) which maximizes the number of usable reads while reducing the sequencing errors of CCS reads. We tested the utility of PacRAT by simulating eight libraries of various read lengths and analyzing two real libraries.

## Methods

PacRAT is a reference-free, highly reliable approach for linking barcodes to variants and improves upon the output of a previously used method, Assembly By PacBio (ABP) (Matreyek *et al*., 2018). ABP assigns barcodes to variants by taking either the highest-frequency sequence associated with each barcode or highest average quality sequence associated with each barcode. Crucially, this assigned variant often retains sequencing errors (most commonly 1 bp indels), depending on the coverage and quality of reads associated with each barcode.

To fix this issue, our program assigns barcodes to variants by aligning CCS reads with the same barcode using multiple sequence alignment (Edgar, 2004). A consensus sequence is then generated from these alignments. If no ambiguous nucleotides are present, this consensus sequence is used as the final, error-corrected sequence. However, if ambiguities in these alignments persist, a second pairwise alignment is performed with EMBOSS Needle (Rice *et al*., 2000) between the highest average quality CCS read and the derived consensus sequence. Sites with ambiguities are resolved by taking the nucleotide sequence from the highest quality read. Thresholds for determining consensus sequences can be specified by the user, with the default requiring that the majority of sequences share the same base at each position.

## Results and Conclusions

To assess the accuracy of PacRAT, we computationally generated eight perfect DMS libraries with a range of sizes (∼800 bp to ∼4000 bp) with 5X barcode coverage. Using SimLoRD, a PacBio SMRT read simulator (Stöcker *et al*., 2016), we simulated CCS reads generated from the DMS libraries with 10X read coverage (Figure 1, Supplementary Figure 1, Supplementary Table 1). PacRAT improved barcode-variant mapping vs. ABP in all simulated libraries (Figure 1, Supplementary Figure 1). For the largest simulated library (4086 bp; 201,120 total barcodes), PacRAT correctly identified the variant for 97% of barcodes, whereas ABP only identified 14% (Figure 1). Given the high accuracy, we tested whether we could reduce sequencing coverage and still obtain good library characterization, which could reduce library sequencing costs and improve throughput. By downsampling our libraries to 5X and 3X read coverage, 72% and 41% of barcodes from the longest simulated library were correctly identified, respectively. In a smaller 2.1 kb library, we observed that 99.7%, 91.5%, and 72.7% of barcodes were correctly identified with PacRAT at 10X, 5X, and 3X coverage, respectively. These results indicate that PacRAT can accommodate reduced sequencing read requirements while maintaining high quality data.

**Fig. 1.**
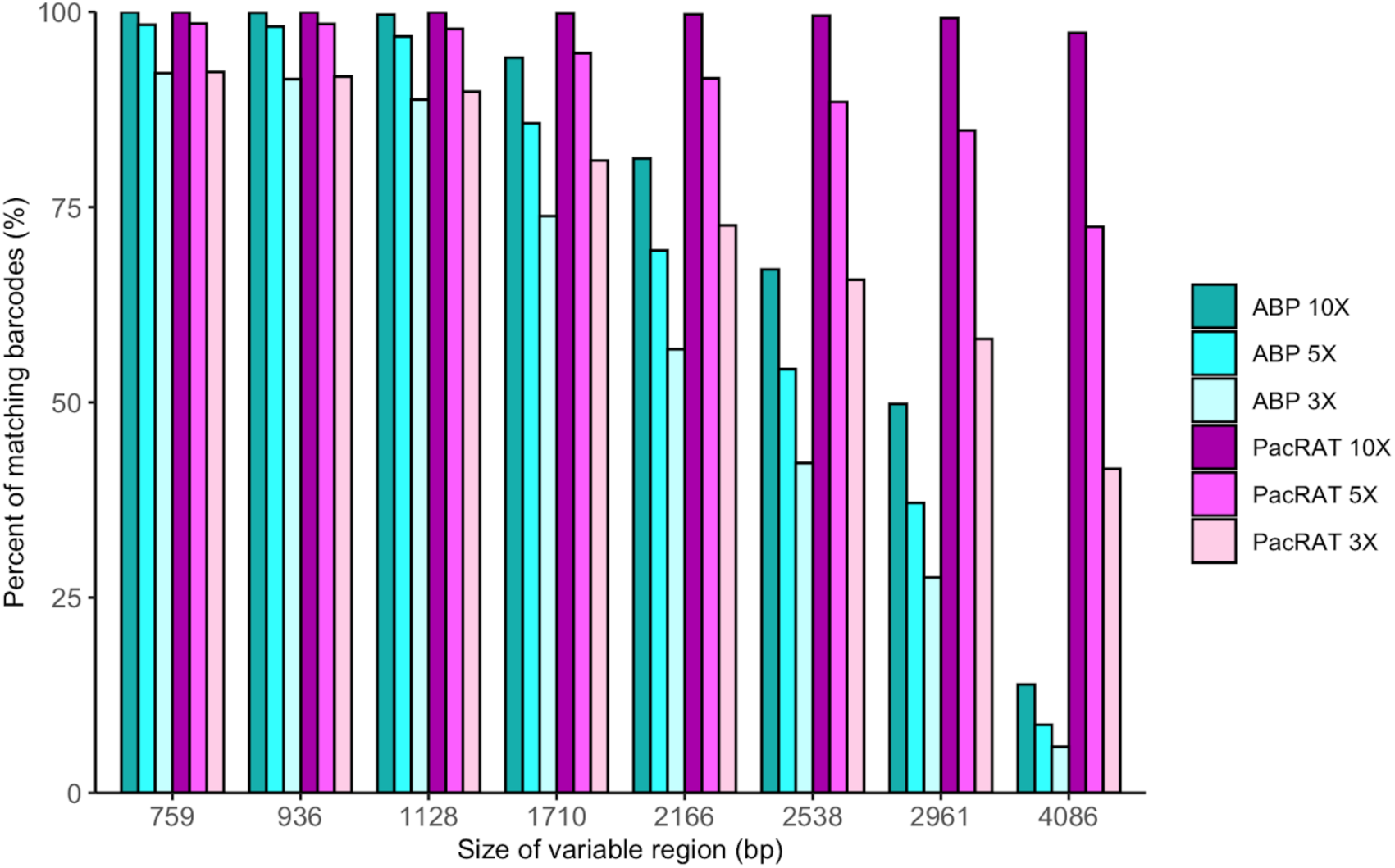
Comparison of correct variant identification between ABP and PacRAT (--cutoff 1, --threshold 0.6) in simulated libraries with different sizes of variable regions. Variant libraries of eight different variable region lengths (759, 936, 1128, 1710, 2166, 2538, 2961, and 4086 bp) were generated using SimLoRD (Stöcker *et al*., 2016) with parameters -xn 8e-03 2.54 5500 -xs 2.78e-03 -0.48 825 48303.073 1.469 chosen to better simulate longer variable regions (Supplementary Table 1) at 10X, 5X, and 3X read coverage.

To verify that our program improves barcode variant mapping with real PacBio reads, we re-analyzed two published variant libraries created for genes *MSH2* (Ollodart *et al*., 2021) and *CYP2C9* (Amorosi *et al*., 2021). The *MSH2* library was generated via DNA synthesis, where specific single amino acid variants were requested from Twist Biosciences. PacRAT decreased the number of unexpected variants (those with synonymous variants, more than one nonsynonymous mutation, and single nonsynonymous mutations not included in the original request) from 16.8% of barcodes analyzed with ABP to 10.5% with PacRAT (Supplementary Table 2). Errors persistent after PacRAT analysis suggest DNA synthesis or sequencing errors.

For the ∼118,000-barcode *CYP2C9* library, which was generated by NNK mutagenesis at each codon via inverse PCR, PacRAT increased the number of barcodes associated with on-target variants from ∼36,100 barcodes to ∼39,800 barcodes, while reducing the number of barcodes associated with indels from ∼66,800 barcodes to ∼61,600 barcodes. Barcodes associated with indels are expected to have low functional scores, but with ABP only, we observed 3,043 barcodes associated with indels that had WT-like scores, likely indicating a variant misclassification. However, PacRAT was able to reduce the number of these misclassifications by ∼50%.

PacRAT improves upon existing PacBio library barcode-variant mapping pipelines by implementing multiple sequence alignments to resolve the variant sequence associated with each barcode. Our method is reference-free and resolves sequence artifacts, commonly indels. While not all reads will map to an expected variant, users can adjust thresholds to determine the ideal parameters for each specific library. Our approach improves data quality and reduces costs of sequencing by utilizing reads that were assigned incorrect sequences or otherwise discarded in previous methods. PacRAT also provides researchers with metrics for estimating the necessary read coverage for different types of libraries.

## Acknowledgements

Thanks to Phil Green for his original suggestion of this approach. We thank Jochen Weile, Atina Cote, and Fritz Roth for the helpful suggestions to improve PacRAT. Thanks to Nick Popp for beta testing PacRAT.

## Supplemental Data

**Supplementary Figure 1.**
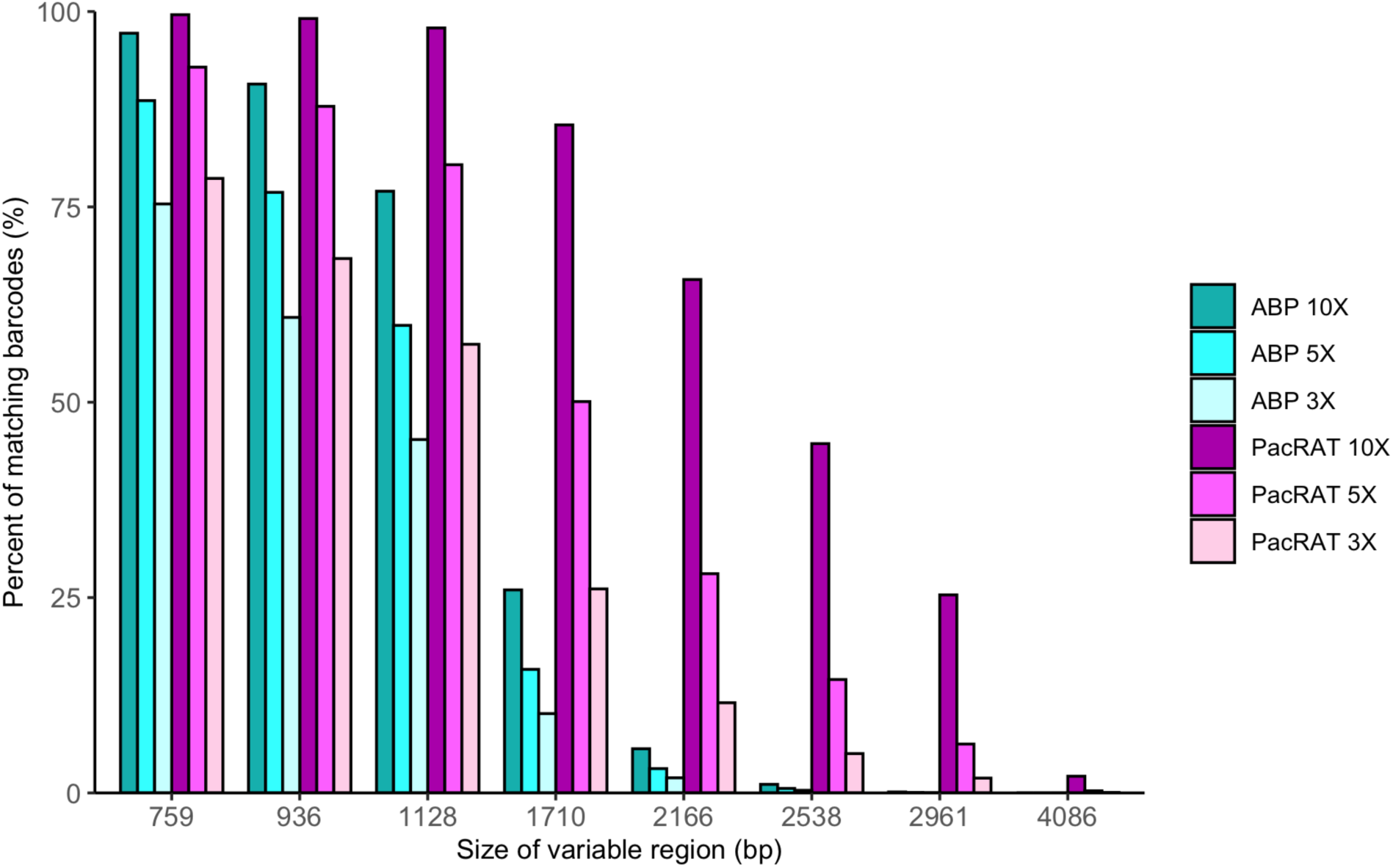
Comparison of correct variant identification between ABP and PacRAT in simulated libraries with default SimLoRD settings. Reads from libraries with eight different variable region lengths (759, 936, 1128, 1710, 2166, 2538, 2961, and 4086 bp) were generated using SimLoRD with default Chi-squared parameters (-xn 1.8923e-03 2.5394e+00 5500 -xs 0.0121 -5.12 675 48303.073 1.469) at 10X, 5X, and 3X read coverage. These default parameters are designed to model P4 chemistry (RS II). The number of correctly identified variants using ABP and PacRAT were compared. PacRAT increased the number of correctly identified variants in libraries sized between 1700-2500 bp, but its effectiveness dropped rapidly after 2500 bp.

**Supplementary Table 1.** Comparison of mean passes through SimLoRD simulated and real libraries. We determined that the average pass numbers produced using SimLoRD’s default Chi-squared parameters (-xn 1.8923e-03 2.5394e+00 5500 -xs 0.0121 -5.12 675 48303.073 1.469) did not accurately reflect those of real PacBio libraries. Thus, we adjusted these parameters (-xn 8e-03 2.54 5500 -xs 2.78e-03 -0.48 825 48303.073 1.469) to better reflect the number of passes found in Sequel II data.

| Gene | length (bp) | mean number of passes | real or simulated library | default or adjusted SimLoRD parameters |
| --- | --- | --- | --- | --- |
| MSH2 | 4675 | 11.55 | real | NA |
| STAG1 | 4086 | 4.34 | simulated | default |
| STAG1 | 4086 | 9.65 | simulated | adjusted |
| CYP2C9 | 1625 | 19.45 | real | NA |
| CEP55 | 1710 | 6.51 | simulated | default |
| CEP55 | 1710 | 14.86 | simulated | adjusted |

**Supplementary Table 2.**
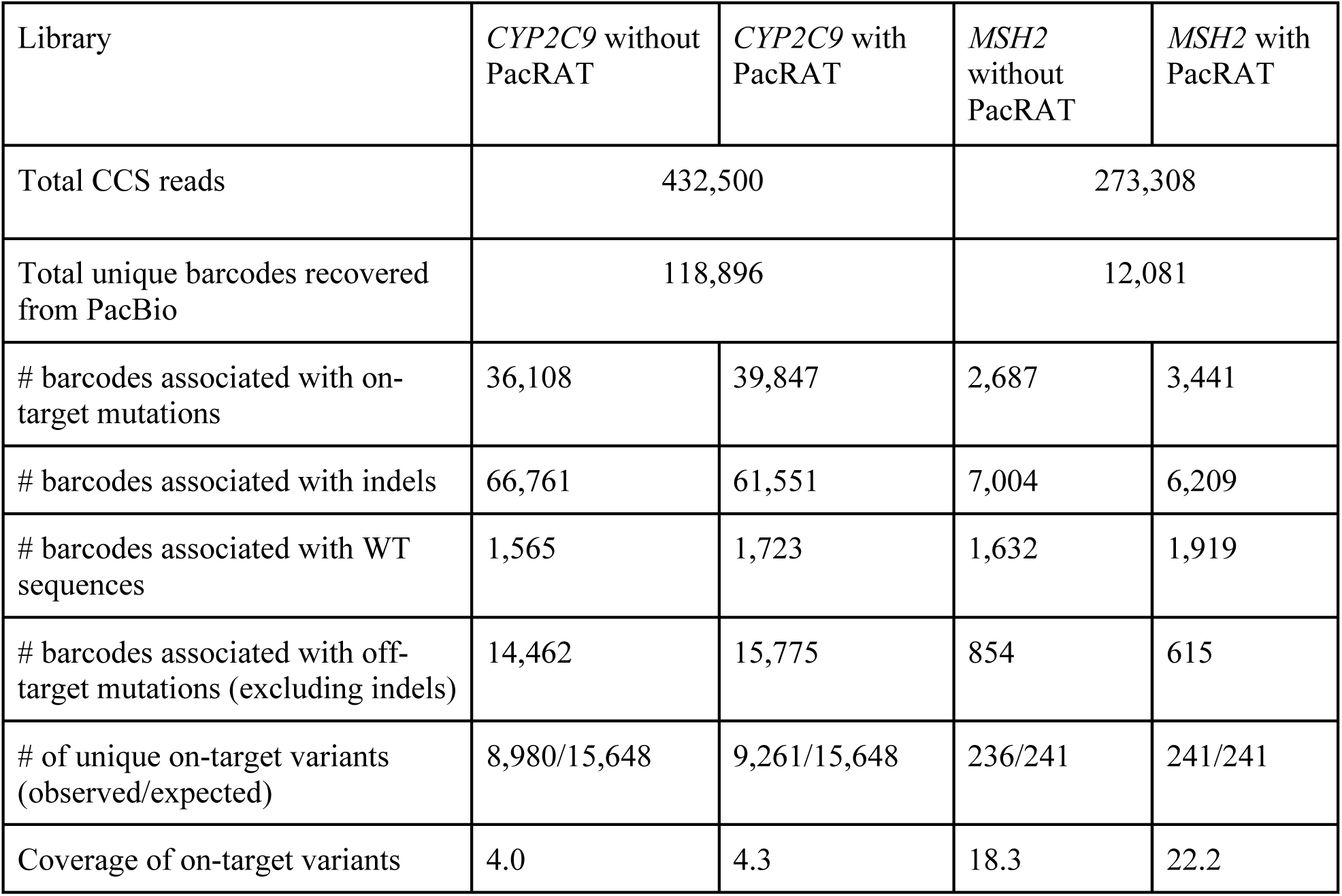
*CYP2C9* and *MSH2* library statistics. The *MSH2* library was synthesized from Twist Biosciences and sequenced on a Sequel II machine (Sequel 3.0) using one flow cell. The *CYP2C9* library is the activity library described in Amorosi et al. 2021, and was sequenced on a Sequel II machine (Sequel 2.1) using two flow cells. For PacRAT analysis, parameters --cutoff 1 and --threshold 0.6 were used.

| Library | *CYP2C9* without PacRAT | *CYP2C9* with PacRAT | *MSH2* without PacRAT | *MSH2* with PacRAT |
| --- | --- | --- | --- | --- |
| Total CCS reads | 432,500 |  | 273,308 |  |
| Total unique barcodes recovered from PacBio | 118,896 |  | 12,081 |  |
| # barcodes associated with on-target mutations | 36,108 | 39,847 | 2,687 | 3,441 |
| # barcodes associated with indels | 66,761 | 61,551 | 7,004 | 6,209 |
| # barcodes associated with WT sequences | 1,565 | 1,723 | 1,632 | 1,919 |
| # barcodes associated with off-target mutations (excluding indels) | 14,462 | 15,775 | 854 | 615 |
| # of unique on-target variants (observed/expected) | 8,980/15,648 | 9,261/15,648 | 236/241 | 241/241 |
| Coverage of on-target variants | 4.0 | 4.3 | 18.3 | 22.2 |

